# Plasticity of circadian and circatidal rhythms in activity and transcriptomic dynamics in a freshwater snail

**DOI:** 10.1101/2023.08.19.553957

**Authors:** Takumi Yokomizo, Yuma Takahashi

**Affiliations:** Graduate School of Science, Chiba University, Chiba 263-8522, Japan

**Keywords:** endogenous rhythm, biological clock, phenotypic plasticity, tidal environment, transcriptome

## Abstract

Organisms have diverse biological clocks synchronized with environmental cycles depending on their habitats. Anticipation of tidal changes has driven the evolution of circatidal rhythms in some marine species. In a freshwater snail, *Semisulcospira reiniana*, individuals in nontidal areas exhibit circadian rhythms, whereas those in tidal areas exhibit both circadian and circatidal rhythms. We investigated whether the circatidal rhythms are genetically determined or induced by environmental cycles. After exposure to a simulated tidal cycle, both tidal and nontidal populations exhibited circatidal rhythms in locomotive activity. Transcriptome analysis revealed that genes with circatidal oscillation increased due to entrainment to the tidal cycle in both populations and dominant rhythmicity was consistent with the environmental cycle. These results suggest plasticity in the endogenous rhythm in both populations. However, circatidal oscillating genes were more abundant in the tidal population than in the nontidal population, suggesting that a greater number of genes are associated with circatidal clocks in the tidal population compared to the nontidal population. Our findings suggest that the plasticity of biological rhythms and genetic changes in the molecular network between clocks and downstream genes may have contributed to the adaptation to the tidal environment in *S. reiniana*.

## Introduction

Organisms have faced the challenge of adapting to periodic variations in environments since ancient times. The rotation and inertial force of Earth, along with the gravitational forces of the sun and moon, generate environmental variations across multiple timescales, including day–night, tidal, lunar, and seasonal cycles. To cope with these fluctuations, organisms possess self-sustained, temperature-compensated timekeeping systems capable of entraining to environmental cues [1–3]. The circadian rhythm, observed in a wide range of organisms, ranging from bacteria [4] to mammals [5–7], allows for adaptation to day–night cycles. Biological clocks enable organisms to anticipate environmental changes and regulate physiological processes with appropriate timing. The coordination between environmental cycles and biological clock parameters, such as phase and period, plays a vital role in enhancing fitness and facilitating adaptation to rhythmic environments [8–10].

Synchronizing biological rhythms with environmental cycles has adaptive significance and can result in variations in endogenous rhythms depending on the circumstances. For instance, studies on the model fungus *Neurospora discreta* have shown that local adaptation to different rhythmic environments, regulated by the circadian clock, leads to habitat-specific endogenous rhythms and influences reproductive success [11]. Variations in endogenous rhythms can arise not only from local adaptation but also from plasticity driven by social roles. Honeybees, for example, exhibit social environment-dependent plasticity in their circadian rhythm within a single population [12,13]. These findings suggest that genetic and/or nongenetic changes in biological rhythms contribute to enhancing individual performance.

Organisms residing in marine or intertidal habitats encounter substantial and complex environmental changes, including fluctuations in water level, salinity, temperature, light, and food availability, driven by daily and tidal cycles. The circatidal rhythm, an approximately 12-h biological rhythm synchronized with tides, is observed in diverse taxa [14–17]. Owing to the critical importance of anticipating tidal changes for marine species, the selective pressure exerted by the tidal cycle can drive the adaptive evolution of circatidal clocks. The relationship between circadian and circatidal rhythms, characterized by different periodicities and zeitgebers, remains controversial. For example, studies on the mangrove cricket suggest that the circatidal timekeeping system operates independently of circadian clocks [18,19]. This dissociation in the circadian and circatidal clocks of mangrove crickets may be attributed to the adaptive evolution of a novel timekeeping system that emerged in their mangrove habitat. Conversely, investigations of various marine species have demonstrated plasticity in rhythm expression within individuals or populations under different environmental conditions [20–22]. Notably, certain circadian clock genes in the Pacific oyster *Crassostrea gigas* exhibit oscillation at tidal frequencies under tidal conditions [23,24]. In the amphipod crustacean *Parhyale hawaiensis*, the core circadian clock gene *Bmal1* is suggested to be essential for circatidal rhythms [25]. These studies give rise to the hypothesis that the circadian and circatidal rhythms may be generated by a single biological timekeeping system, with circatidal clocks sharing some molecular components with circadian clocks. The expression of circatidal rhythms and successful adaptation to tidal environments likely involve a combination of plasticity and genetic changes.

Rivers present complex rhythmic environments due to diurnal and tidal cycles. Although a river follows a 24-h light–dark (LD) cycle throughout its course, downstream areas experience an environmental cycle with a period of 12.4 hours due to the tidal cycle. To investigate variations in endogenous rhythm across habitats with distinct environmental cycles, we focused on the freshwater snail *Semisulcospira reiniana*, which inhabits both nontidal and tidal areas of rivers. Our previous study revealed that snails in nontidal areas exhibit circadian rhythms, whereas those in tidal areas exhibit both circadian and circatidal rhythms [26]. However, it remains unclear whether the differential rhythmicity arises from evolutionary differentiation or plasticity in response to environmental cycles. In the present study, we evaluated genetic and nongenetic changes in the endogenous rhythm of snails living in habitats characterized by heterogeneous environmental cycles. We exposed snails to a simulated tidal environment in the laboratory and examined the activity and transcriptome rhythms of individuals from both tidal and nontidal populations.

## Materials and Methods

### Species and sampling sites

*Semisulcospira reiniana*, a common freshwater snail species found in Japan, was collected from nontidal (3–5 m above sea level; 35° 15′ 15″ N, 136° 41′ 09″ E) and tidal areas (<1 m above sea level; 35° 08′ 49″ N, 136° 40′ 37″ E) of the Kiso River in June and November 2021. The two populations were approximately 20 km apart along the river. The collected snails were acclimated in the laboratory under 23°C and 12–12-h LD conditions in freshwater for at least one month.

### Entrainment to the simulated tidal cycle

After acclimation, snails from the tidal and nontidal populations were divided into control and treatment groups. The treatment group individuals were exposed to a simulated tidal cycle by raising and lowering the water level in a water tank (60 × 30 × 45 cm; Fig. S1). This tidal simulation was conducted twice in the laboratory under 23°C and 12–12-h LD (12L12D) conditions. Containers (φ7 × 9 cm) with water vents were placed 32 cm from the bottom of the tank. Four snails were placed in each container. For the high tide condition, the water level was filled to a depth of 39 cm, submerging the snails completely. To simulate the low tide condition, the water level was dropped to a depth of 24 cm, exposing the snails completely to air. Water supply and draining were controlled by timers. In the first simulation, water supply switched to drainage at 03:00 and 15:00, and drainage switched to supply at 09:00 and 21:00. In the second simulation, water supply switched to drainage at 05:00 and 17:00, and drainage switched to supply at 11:00 and 23:00. The entrainment treatment lasted for four weeks. Snails from the first tidal simulation were used for behavioural observations, whereas those from the second simulation were used for gene expression analysis. Individuals in the control group were kept in freshwater without water level oscillation under 12L12D conditions.

### Behavioural observation

To examine the effect of 12-h water level oscillation on the endogenous activity rhythm of *S. reiniana*, individuals from the control and treatment groups in both nontidal and tidal populations were observed under constant laboratory conditions. Eight snails per population were individually placed in containers (13 × 11.7 cm, 4.7 cm in height) filled with freshwater. The snails were observed under constant darkness (DD) at a temperature of 23°C in the laboratory for 96 and 84 h in the control and treatment groups, respectively. A camera malfunction resulted in a shorter observation period for the treatment group compared with the control group. Water circulation was maintained using a pump to keep the water clean. Images were captured every 30 s using a model 400-CAM061 camera (Sanwa Direct, Japan) and were used to create time-lapse movies. UMATracker tracking software [27] was used to track the positions of snails. After correcting major tracking errors, the coordinates of the snails were averaged every three frames to minimize the influence of small deviations. The total locomotion distance travelled in 1 h was calculated using the snail trajectories.

### RNA sampling

To investigate the effect of a 12-h water level oscillation on the endogenous gene expression rhythm of *S. reiniana*, time-course RNA sequencing (RNA-seq) was conducted using individuals from the control and treatment groups in both nontidal and tidal populations. Snails were preserved in containers without water under the DD condition at a constant temperature (23°C) for tissue collection. Starting at subjective high tide, three individuals per population were dissected every 3.1 h for a total of 49.6 h (17 sampling time points) in both control and treatment groups. The epidermis of each individual was preserved in 750 μL of RNA*later* Stabilization Solution (Invitrogen). The samples were stored at 4°C for approximately 3 h and then stored at −80°C until total RNA extraction.

### RNA extraction and sequencing

Total RNA was extracted using the Maxwell 16 LEV Plant RNA Kit with the Maxwell 16 Research Instrument (Promega, USA) following the manufacturer’s instructions. Electrophoresis was performed on a 1% agarose gel to assess RNA degradation. RNA concentrations were estimated using the Qubit 2.0 fluorometer (Invitrogen), and RNA purity was assessed using the NanoDrop Lite Spectrophotometer (Thermo Scientific). Total RNA from three individuals in the same population at each sampling time point was pooled in equal amounts for pooled RNA-seq. A cDNA library was constructed using the TruSeq RNA Sample Prep Kit, and paired-end (150 bp) pooled RNA-seq was performed on the Illumina NovaSeq6000 platform. Adaptor sequences and low-quality reads were removed using Trimmomatic (version 0.38) [28], and FastQC (version 0.11.8; http://www.bioinformatics.babraham.ac.uk/projects/fastqc/) was used for quality control. The remaining high-quality reads from all samples were used for *de novo* assembly via Trinity software (version 2.9.1) [29]. To estimate gene expression levels, reads from each sample were mapped to the reference transcripts using RSEM software (version 1.3.0) [30] to obtain the transcripts per million (TPM) for each gene. The reference transcripts were used to create supertranscripts, and a BLAST search was performed against all protein sequences of *C. gigas* [31] to identify homologs for each gene with the best hit and an e-value <0.0001. Genes showing circatidal oscillation were analyzed based on the identified homologs.

### Statistical analyses

The activity rhythm of individuals in the control and treatment groups was examined using Lomb–Scargle periodogram analysis via ActogramJ, a software package based on ImageJ for chronobiological data analysis and visualization [32]. The mean locomotor distance of surviving individuals was used for rhythm detection. Owing to the limited movement of individuals in the treatment group at the end of the observation, activity data from 72 h after the onset were used for the periodogram analysis. For transcriptome analysis, genes with low expression (average TPM ≤ 1) or small oscillation amplitude (peak/trough ≤ 1.3) were excluded. Probabilistic principal component analysis (PCA) was performed to summarize the gene expression pattern of each sample. After removing an outlier sample, Tukey’s honestly significant difference (HSD) test was conducted to identify differences in expression patterns between populations or entrainment treatments. Gene expression rhythmicity was analyzed using RAIN [33], a nonparametric algorithm for identifying rhythmic components in large biological datasets. Circatidal (12.4 ± 3.1 h) and circadian (24.8 ± 3.1 h) oscillating genes (*p* < 0.01) were identified. A differential rhythmicity score (*S*_DR_) was calculated for each gene to assess the differential rhythmicity of gene expression between the control and treatment groups. Based on periodicity analysis performed via RAIN, genes with a P-value of circatidal periodicity <1 were used for the calculation of *S*_DR_, which was defined as follows:

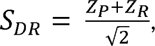

where *Z_P_* and *Z_R_* represent the *Z*-scores for changes in periodicity and amplitude between groups, respectively [34]. Changes in periodicity were calculated as log (P-value of the control group) − log (P-value of the treatment group), whereas changes in amplitude were calculated as log_2_ (amplitude of the treatment group / amplitude of the control group). The amplitude of each gene expression was defined as the difference between the maximum and minimum TPM values. The P-value for *S*_DR_ was computed using a Gaussian distribution fitted to the empirical distribution, and it was adjusted using the Benjamini & Hochberg procedure for multiple testing. To assess the relative significance of circadian and circatidal rhythms in the transcriptome, the log-scaled ratio of the P-value for circadian periodicity to that for circatidal periodicity was calculated for each gene. Functional enrichment analysis of biological processes in circatidal oscillating genes or differential rhythmic genes based on *S*_DR_ was performed using Kyoto Encyclopedia of Genes and Genomes (KEGG) pathway over-representation analysis and KEGG gene set enrichment analysis (KEGG-GSEA) conducted through clusterProfiler [35]. Circatidal oscillating genes identified via RAIN in the treatment group were used as the test set for KEGG pathway over-representation analysis. The gene list for GSEA was generated based on *S*_DR_. All statistical analyses were conducted using R version 4.2.2.

## Results

### Activity rhythm

Snails exhibited rhythmic activity patterns under DD conditions in the laboratory, with a gradual decrease in activity observed in the treatment group (Figs. 1 and 2). In the control group, individuals from the nontidal and tidal populations exhibited a circadian activity rhythm with periods of 21.6 h and 22.5 h, respectively, under laboratory DD conditions (Figs. 1a, b). The tidal population did not exhibit the circatidal rhythm, indicating synchronization with the laboratory 12L12D condition. In contrast, the treatment group exhibited a circatidal activity rhythm with periods of 12.4 h and 11.9 h in the nontidal and tidal populations, respectively, under laboratory DD conditions (Figs. 2a, b). This indicates entrainment to the simulated tidal cycle, with the activity peak preceding the expected high tide peak.

**Figure 1.**
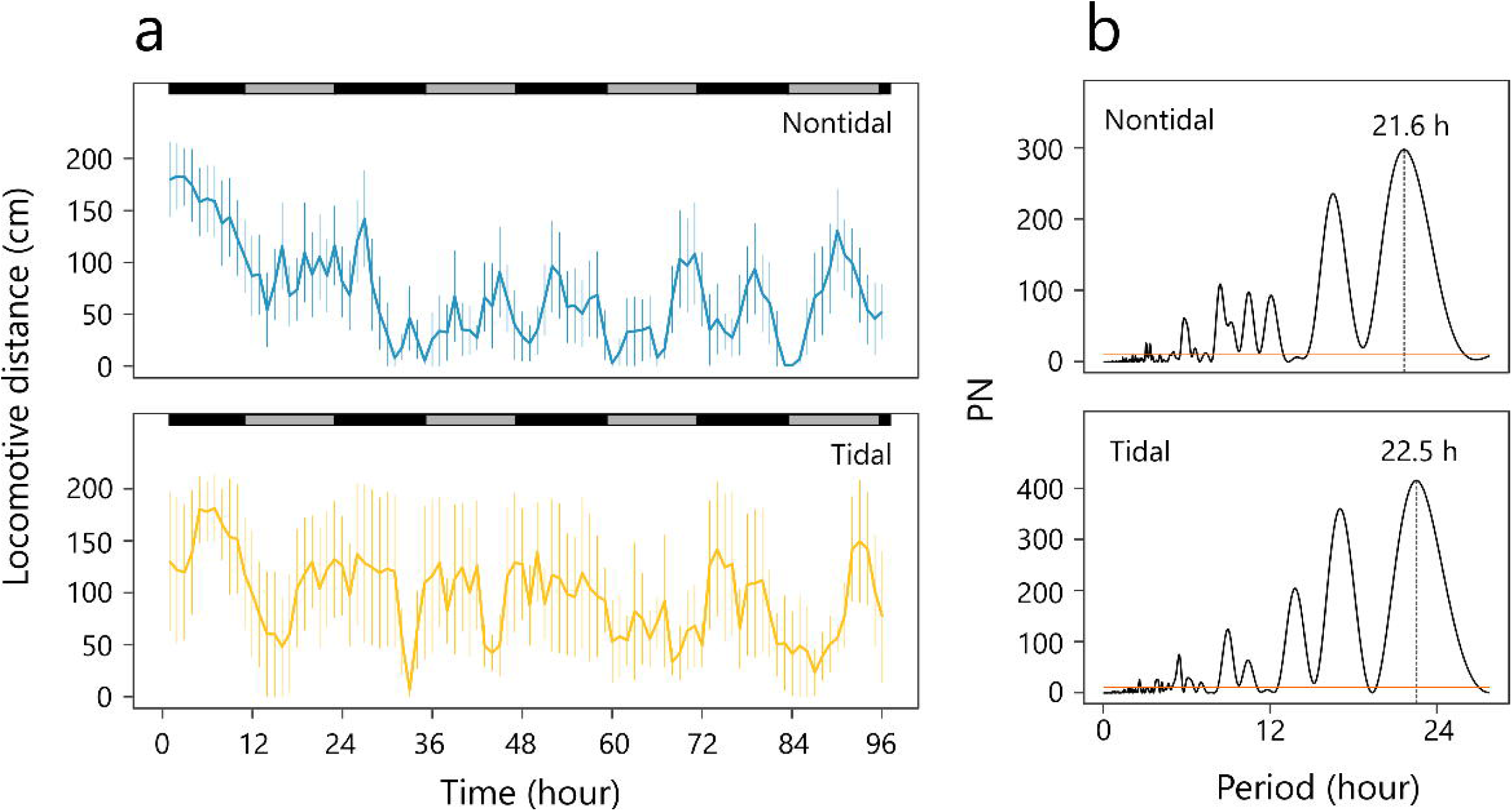
Activity patterns of individuals in the nontidal and tidal populations of the control group under laboratory DD conditions. (a) Locomotor distance of the nontidal and tidal populations over 96 h. Error bars represent the standard error of the mean (SEM). Subjective day and night are indicated by grey and black bars above the patterns, respectively. (b) Lomb–Scargle periodogram analysis showing peaks in the circadian range.

**Figure 2.**
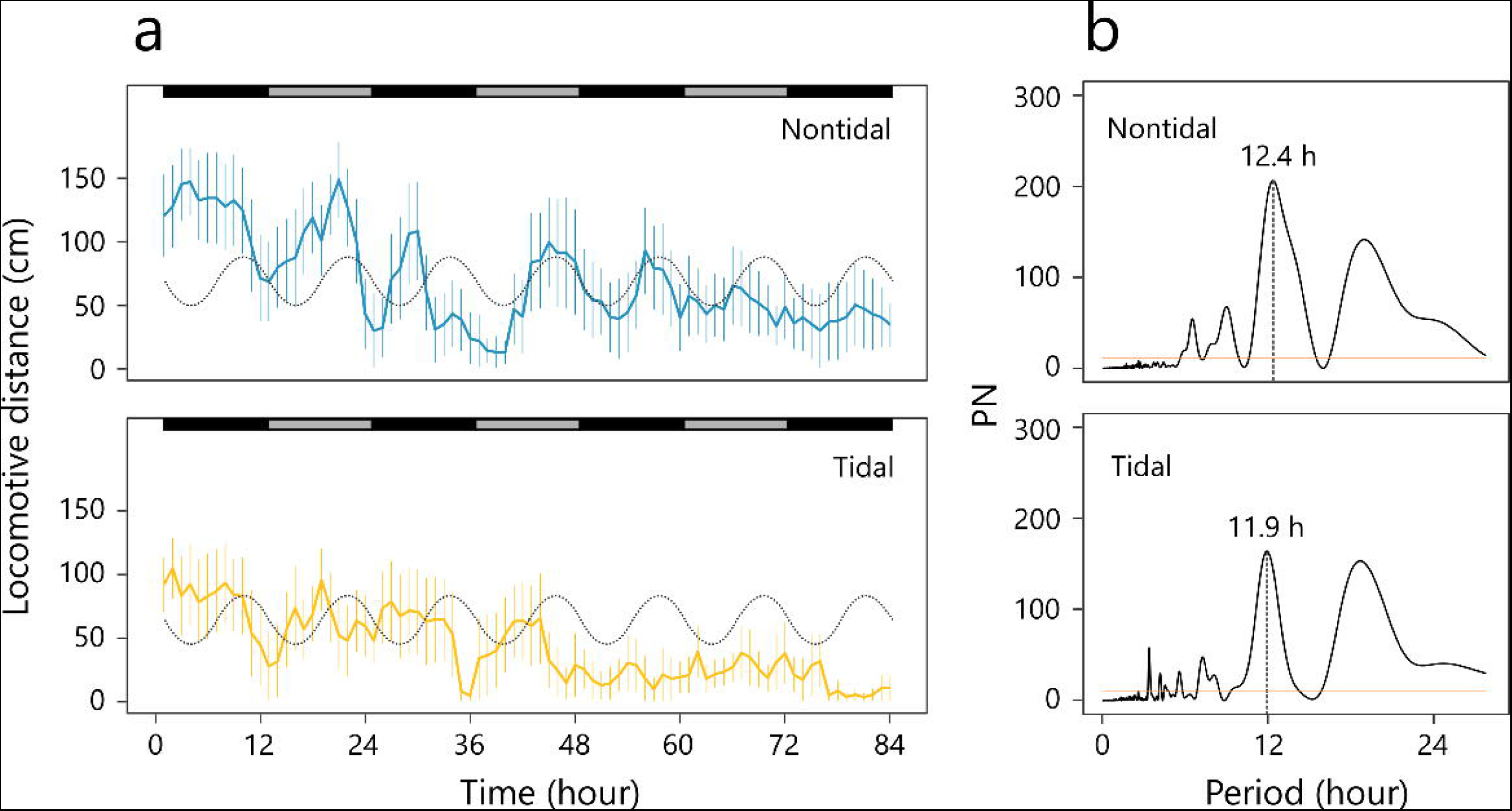
Activity patterns of individuals in the nontidal and tidal populations of the treatment group under laboratory DD conditions. (a) Locomotor distance of the nontidal and tidal populations over 84 h. Error bars represent the SEM. Subjective day and night are indicated by grey and black bars above the patterns, respectively. The expected tide level in the laboratory is shown as a grey dotted line. (b) Lomb–Scargle periodogram analysis showing peaks in the circatidal range.

### Transcriptome rhythm

In total, we obtained 1,578,392 contigs with a mean length of 513 bp. Among the 954 metazoan core gene orthologs, we identified 948 (99.3%). We obtained 908,817 supertranscripts containing 58,952 contigs annotated against all protein sequences of *C. gigas*. In the tidal population, 13,929 and 14,508 genes in the control group and treatment group, respectively, met the criteria of average TPM value >0 and oscillation amplitude (peak/trough) ≥1.3. In the nontidal population, the corresponding numbers were 13,448 and 13,997 genes in the control group and treatment group, respectively. Based on probabilistic PCA, one sample in the control group of the nontidal population showed a greatly different expression pattern compared with the other samples (Fig. S2) and was excluded from subsequent analysis. Probabilistic PCA of the remaining samples indicated significant differences in expression patterns between populations but not between the control and treatment groups (Fig. 3a; Table 1). In the nontidal population, we identified 190 (1.4%) and 292 (2.1%) circatidal (12.4 ± 3.1 h) oscillating genes in the control group and treatment group, respectively (Figs. 3b–d). In the tidal population, the corresponding numbers were 271 (1.9%) and 340 (2.3%) circatidal oscillating genes in the control group and treatment group, respectively (Figs. 3b–d). The proportion of genes oscillating in the circatidal period increased in both populations with entrainment to the simulated tidal cycle. However, the abundance of oscillating genes was higher in the tidal population compared with the nontidal population, regardless of the control or treatment group. We also detected several oscillating genes with a circadian period. In the nontidal population, 343 (2.6%) and 328 (2.3%) oscillating genes were found in the control and treatment groups, respectively (Figs. 3b–d). In the tidal population, the corresponding numbers were 420 (3.0%) and 301 (2.1%) in the control and treatment groups, respectively (Figs. 3b–d). Unlike circatidal oscillating genes, the proportion of genes oscillating in the circadian period decreased in both populations with entrainment to the simulated tidal cycle. No circadian clock genes showed circadian or circatidal rhythmicity in the control and treatment groups in both populations (Figs. S3 and S4).

**Figure 3.**
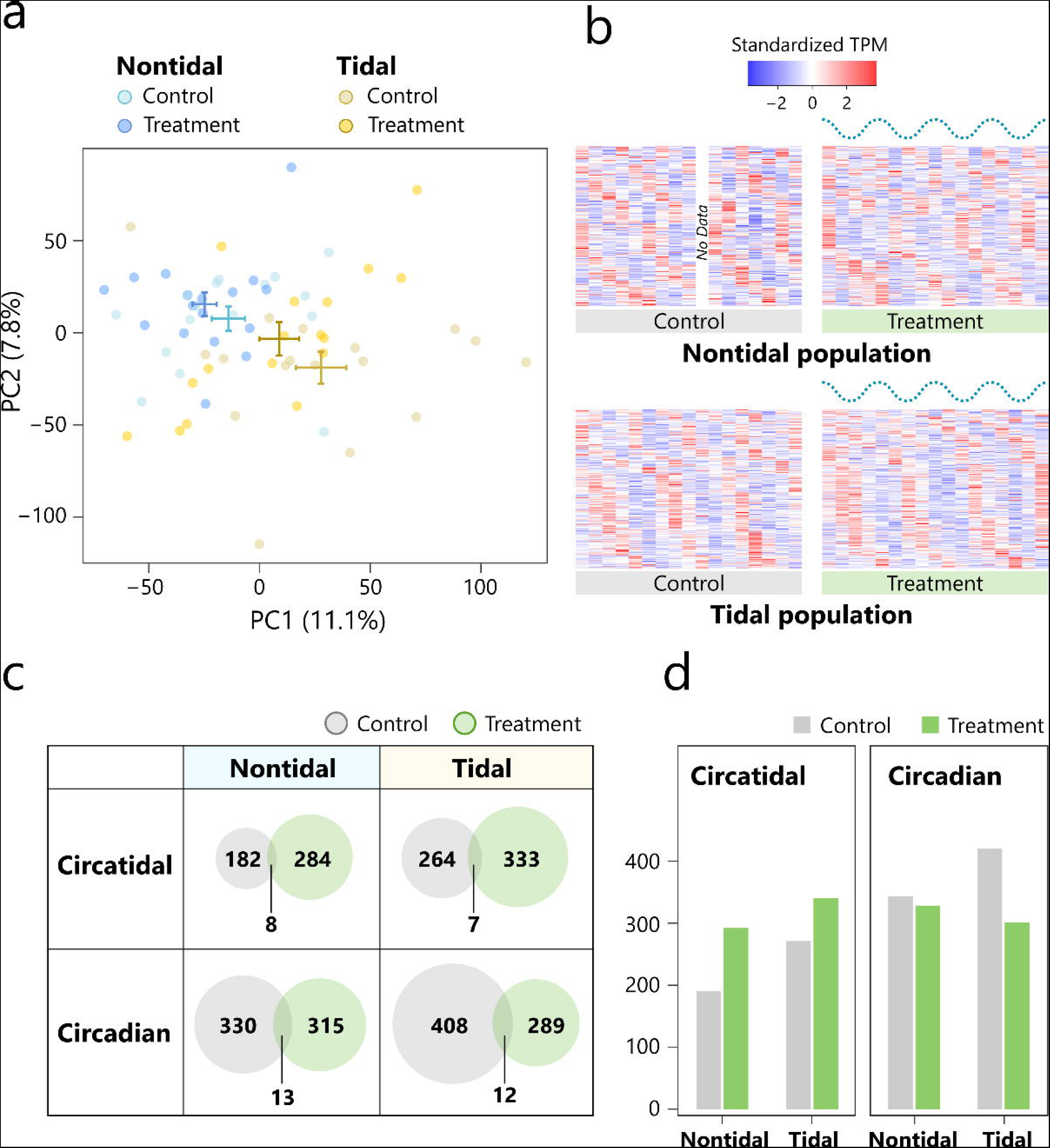
Gene expression patterns and rhythms of individuals in the nontidal and tidal populations under laboratory DD conditions. Control samples were kept under LD conditions without the tidal cycle. Treatment samples were exposed to the simulated tidal cycle for four weeks. (a) Probabilistic PCA of the expression of all genes after filtering in the nontidal and tidal populations. (b) Heatmaps of standardized expression patterns of rhythmic transcripts with circatidal periods in the nontidal and tidal individuals under DD conditions detected via RAIN. The dotted wavy lines above the heatmaps of treatment samples represent the simulated tide level. (c) Venn diagrams detailing the number of circatidal and circadian oscillating genes detected in the nontidal and tidal populations. (d) Bar plot showing the number of circatidal and circadian oscillating genes detected in the nontidal and tidal populations.

**Table 1.** Tukey’s HSD test of differences in gene expression patterns based on PC1.

|  | **Mean difference** | **P-value** |
| --- | --- | --- |
| Nontidal vs. Tidal | 37.67 | $4.7 \times 10^{-5}$ |
| Control vs. Treatment | -14.80 | 0.09 |
| Nontidal Control vs. Nontidal Treatment | -10.74 | 0.82 |
| Nontidal Control vs. Tidal Control | 41.51 | 0.0067 |
| Nontidal Control vs. Tidal Treatment | 22.77 | 0.26 |
| Nontidal Treatment vs. Tidal Control | -52.25 | $3.2 \times 10^{-4}$ |
| Nontidal Treatment vs. Tidal Treatment | 33.51 | 0.036 |
| Tidal Control vs. Tidal Treatment | -18.74 | 0.41 |

Differential rhythmicity analysis revealed changes in transcriptome rhythmicity between the control and treatment groups (Fig. S5). Genes in the first quadrant exhibited clearer periodicity and larger amplitude in the treatment group compared with the control group, indicating increased circatidal rhythmicity. In the nontidal population, 13 genes showed significantly increased circatidal rhythmicity in the treatment group (FDR < 0.05; Table S1). In the tidal population, 16 genes showed significantly increased circatidal rhythmicity in the treatment group (FDR < 0.05; Table S2). Although most transcripts did not show significant differential rhythmicity in expression patterns in the tidal and nontidal populations, the log-scaled ratio of the P-value for circadian periodicity to that for circatidal periodicity changed with entrainment to the tidal cycle (Fig. 4). In both populations, the log-scaled P-value ratio was negative in the control groups, indicating dominance of the circadian rhythm over the circatidal rhythm under LD conditions without a tidal cycle. However, the ratio was positive in the treatment groups, indicating dominance of the circatidal rhythm over the circadian rhythm after entrainment to the tidal cycle. KEGG pathway over-representation analysis revealed that circatidal oscillating genes in the treatment group of the tidal population were enriched for some pathways, including “Aminoacyl-tRNA biosynthesis” (KO: crg00970) and “Histidine metabolism” (KO: crg00340) (Fig. 5a). KEGG-GSEA based on *S*_DR_ revealed enrichment of the biological process “Ribosome” (KO: crg03010) for genes with increased circatidal rhythmicity in the tidal population (Fig. 5b). Based on both KEGG pathway over-representation analysis and KEGG-GSEA, no enriched pathways were detected in the nontidal population.

**Figure 4.**
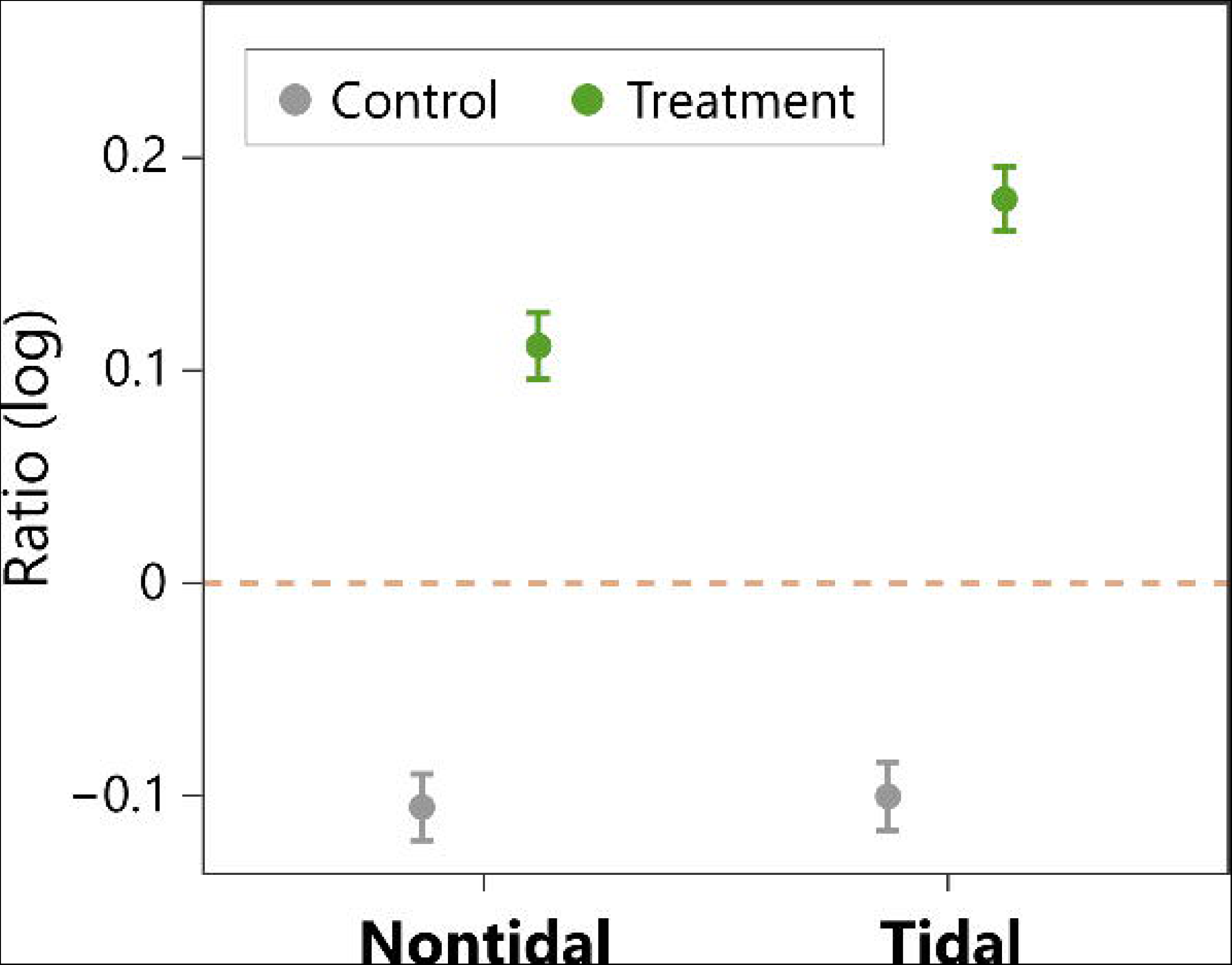
The dominant rhythmicity of the transcriptome in the control and treatment groups of the nontidal and tidal populations evaluated by log-scaled ratio of the P-value of circadian periodicity to that of circatidal periodicity in the transcriptome. The dashed line represents the same P-values for circadian and circatidal periodicity. Error bars represent the SEM.

**Figure 5.**
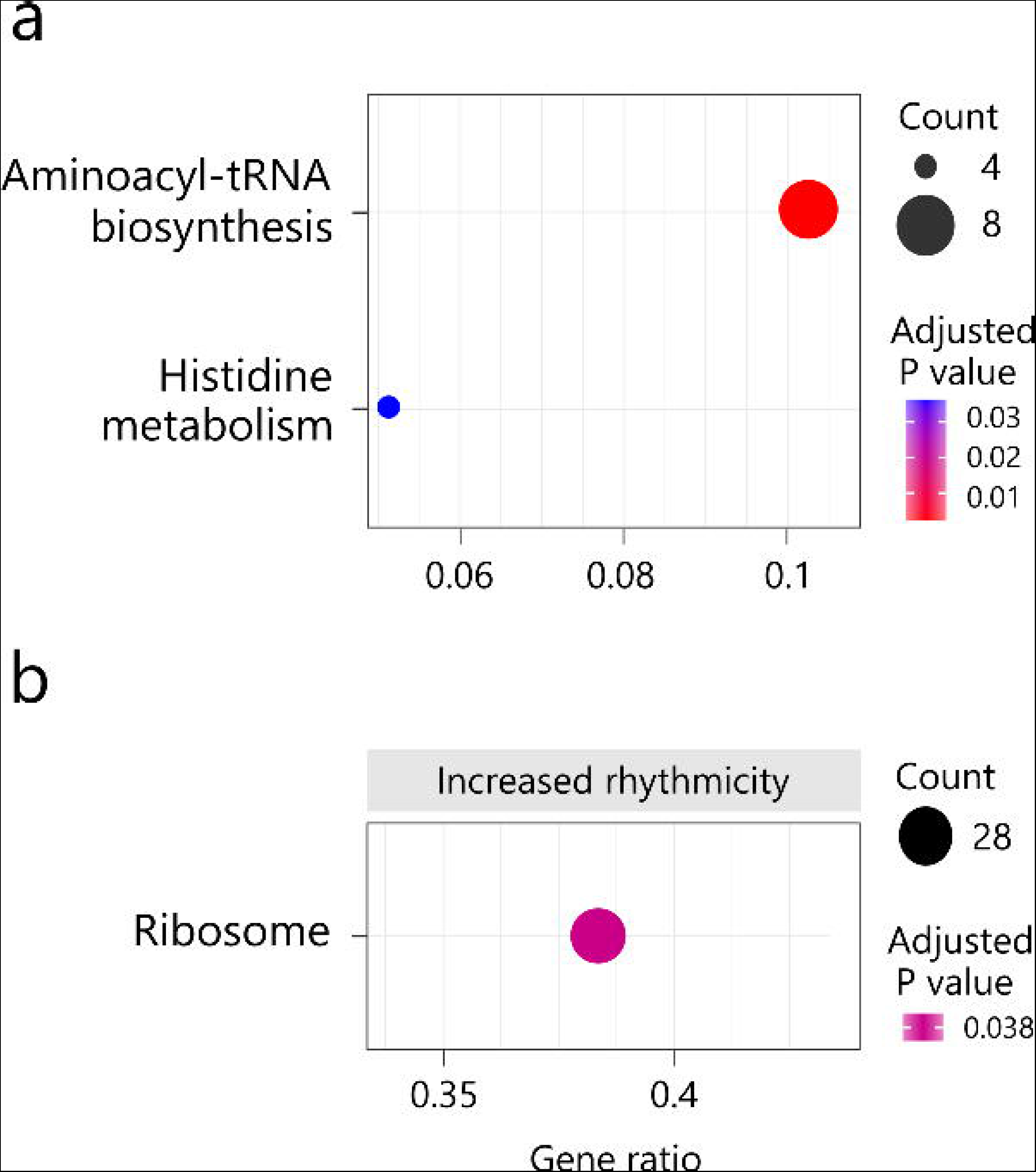
KEGG pathway analysis of the tidal population. (a) KEGG pathway over-representation analysis of the tidal population. Gene ratio refers to the ratio of the number of differentially expressed genes (DEGs) to the total number of genes in a specific pathway. The size of the circles represents the count of DEGs. Circles are coloured based on the adjusted P-value. (b) KEGG-GSEA of the tidal population. Gene ratio refers to the ratio of the number of core enriched genes to the total number of genes in a specific pathway. The size of the circles represents the count of the core enriched genes. No pathways involved in decreased circatidal rhythmicity were detected.

## Discussion

The coordination of biological processes with environmental cycles is regulated by biological clocks, such as the circadian clock, which enhances fitness in organisms in rhythmic environments [36]. Marine and intertidal habitats, characterized by the tidal cycle, experience complex and drastic environmental variations, unlike terrestrial and inland water habitats. The freshwater snail *S*. *reiniana*, found in tidal areas, exhibits the circatidal rhythm, which is believed to be an adaptation to the tidal environment [26]. In the present study, we investigated whether the differential rhythmicity in *S. reiniana* between tidal and nontidal populations is a result of evolutionary differentiation or plastic expression of rhythms depending on dominant environmental cycles. We demonstrated that even individuals in the nontidal population can exhibit the circatidal rhythm through entrainment to the tidal cycle. However, our results regarding the transcriptome rhythm suggest that more genes or pathways are influenced by the tidal cycle in the tidal population compared with the nontidal population.

We first examined the influence of the simulated tidal cycle on the activity rhythm under constant conditions in each population. The free-running period of individuals in the control group under DD conditions was shorter than 24 h, regardless of the population. This short circadian period aligns with Aschoff’s rule, which predicts that diurnal animals have a free-running period longer than 24 h and nocturnal animals, such as *S. reiniana* [37], have a free-running period shorter than 24 h [38]. Although individuals in the control group exhibited a circadian activity rhythm, the periodicity was not distinct. The lack of additional circadian zeitgebers, such as daily temperature cycles, under the 24-h light condition in the laboratory may have contributed to noise or ultradian rhythms. The activity rhythms of individuals were markedly altered by exposure to the simulated tidal cycle. Individuals in the treatment group exhibited a circatidal activity rhythm regardless of the population, indicating that the circatidal rhythm was plastically expressed in response to the tidal cycle, even in individuals from the nontidal population. Our results also suggest that water level fluctuations serve as a zeitgeber for the circatidal rhythm in *S. reiniana*. The peak of the activity rhythm slightly differed from that of snails under a natural tidal condition. Snails captured from tidal areas exhibit a rhythmic activity pattern with a peak that coincides with high tide [26]. However, individuals in the treatment group in the present study exhibited peaks preceding the expected high tide. This discrepancy may be attributed to the earlier arrival of high tide compared with the programmed switching time, as the water supply rate was adjusted to reach high tide in slightly under 6 h. Periodogram analysis also detected peaks around 19 h in the activity rhythms of individuals in the treatment group. The origin of this periodicity, which could result from a shortened circadian rhythm or a combination of circadian and circatidal rhythms, remains unidentified.

Based on probabilistic PCA, we observed distinguishable gene expression patterns between populations rather than between the control and treatment groups, indicating minimal differentiation in transcriptome expression between populations. Focusing on rhythmic transcripts, we found an increase in the number of circatidal oscillating genes in both populations following entrainment to the tidal cycle, accompanied by a decrease in the number of circadian oscillating genes. These results suggest that the circatidal rhythm is plastically expressed in both populations, and the reduction in circadian oscillating genes supports the transition from the circadian to the circatidal rhythm under a single biological timekeeping system. Additionally, we identified a greater number of circatidal oscillating genes in the tidal population compared with the nontidal population, implying that adaptation to the tidal environment may contribute to the increase in circatidal clock–controlled genes in individuals from the tidal population. We also detected hundreds of circatidal oscillating genes in the control group of nontidal populations. The expression rhythm with a period of 12.4 h in these genes might arise from the harmonics of circadian clocks, as observed in mammals [39–41]. Although we identified several circadian clock genes, none of these genes exhibited circadian rhythmicity in their expression. This finding is consistent with previous studies that reported the absence of circadian rhythmic expression in clock genes in several intertidal organisms [22,42,43]. The interference of circatidal rhythms with the rhythmic expression of circadian clock genes and weak peripheral clocks in the epidermis or asynchronization between tissues in the samples could potentially explain the lack of rhythmic expression in these genes.

To assess the extent to which the tidal cycle influenced circatidal rhythmicity in the transcriptome, we calculated the *S*_DR_ for each gene in both populations. The *S*_DR_ distribution in the tidal population was similar to that in the nontidal population, indicating comparable changes in circatidal rhythmicity. The number of genes showing a significant increase in circatidal rhythmicity did not differ substantially between populations. These results suggest that the degree of changes in circatidal rhythmicity in the transcriptome is similar in individuals exposed to the tidal cycle, regardless of population. Although the majority of genes did not exhibit significant changes in the strength of circatidal rhythmicity between the control and treatment groups, we identified 13 and 16 genes with significantly increased rhythmicity in the treatment group of the nontidal and tidal populations, respectively. These genes are likely entrained by the simulated tidal cycle, leading to enhanced circatidal rhythmicity. Additionally, we calculated the ratio of P-values for circadian periodicity to circatidal periodicity for each gene. Comparing the ratios between the control and treatment groups revealed that the dominant rhythmicity of the transcriptome aligns with the environmental cycle.

We conducted KEGG over-representation analysis and KEGG-GSEA to identify enriched biological processes among the circatidal oscillating genes and differential rhythmic genes. In the tidal population, we detected three pathways, whereas no pathways were identified in the nontidal population. Considering the larger number of circatidal oscillating genes in the tidal population, it is likely that individuals in the tidal population possess a higher abundance of circatidal clock–controlled genes and biological processes regulated by circatidal clocks compared with individuals in the nontidal population. The enrichment of the “Aminoacyl-tRNA biosynthesis” (KO: crg00970) and “Ribosome” (KO: crg03010) pathways in the circatidal oscillating genes of the treatment group in the tidal population suggests the involvement of circatidal rhythms in biological processes related to translation, as aminoacyl-tRNA is responsible for delivering amino acids for mRNA-guided protein synthesis at the ribosome. Furthermore, the detection of the “Histidine metabolism” (KO: crg00340) pathway suggests a potential association between proteins containing histidine and the response to the tidal cycle.

Our findings on the activity rhythm demonstrate the plasticity of the endogenous rhythm in individuals from both tidal and nontidal populations. The genetic basis of the circatidal rhythm may be conserved in the genome, with the tidal zeitgeber triggering the expression of the circatidal rhythm in individuals from the nontidal population. The plasticity of the endogenous rhythm likely played a role in the adaptation of *S. reiniana* to tidal areas in rivers. We have previously revealed minimal genetic differentiation between the two populations [26], supporting the contribution of plasticity to range expansion. By investigating other rivers where *S. reiniana* is limited to nontidal areas, we can further explore the relationship between the plasticity of the endogenous rhythm and the establishment in tidal areas. Although our findings suggest the existence of a genetic basis for the circatidal rhythm in individuals from the nontidal population, it remains unclear whether they possess independent circatidal clocks alongside circadian clocks or a single timekeeping system governing both circadian and circatidal rhythms. The latter hypothesis is gaining support, particularly in studies on biological clocks in marine molluscs [22,24,44]. Further investigations are needed to elucidate the genetic mechanisms underlying circatidal clocks in freshwater snails. However, the absence or decrease of circadian rhythm in activity and gene expression after entrainment to the tidal cycle supports the notion that circadian and circatidal rhythms are generated by changes in the periodicity of a single biological clock in response to environmental cycles. Unravelling the molecular relationship between circadian and circatidal clocks would contribute to testing the debated hypotheses on the mechanism of circatidal oscillation generation [45–47].

Our study indicates that individuals in the nontidal population also possess the circatidal clock and express the circatidal rhythm when exposed to the tidal cycle. However, transcriptome analysis suggests that individuals in the tidal population exhibit increased regulation through the circatidal clock. Overall, the plasticity of the endogenous rhythm and subsequent genetic changes in a molecular network involving a small number of genes may contribute to adaptation to the tidal environment. Investigating the parallel evolution of biological clocks using individuals from independent rivers would provide a comprehensive understanding of the mechanisms underlying adaptation to tidal environments, including both genetic and nongenetic changes in biological clocks.

## Acknowledgements

This work was supported by research grants JSPS KAKENHI Grant Numbers 17H03729 and 20K06822, Fujiwara Natural History Public Interest Incorporated Foundation and Research Institute of Marine Invertebrates to Y.T., and Grant-in-Aid for JSPS Fellows Grant Number 21J20682 to T.Y. Computations were partially performed on the NIG supercomputer at ROIS National Institute of Genetics.

## Data Accessibility

All raw transcriptome read data were deposited in the DDBJ Sequenced Read Archive under accession numbers SAMD00632165–SAMD00632232. The activity and transcriptome data are available in the Figshare (doi: 10.6084/m9.figshare.23672424 and doi: 10.6084/m9.figshare.23672691, respectively).

## Author Contributions

T.Y. and Y.T. designed the experiments. T.Y. performed the experiments and conducted the data analysis. T.Y. drafted the manuscript. T.Y. and Y.T. edited the manuscript.

## References

1. Dunlap JC. 1999 Molecular bases for circadian clocks. Cell 96, 271–290. (doi:10.1016/S0092-8674(00)80566-8)

2. Bell-Pedersen D, Cassone VM, Earnest DJ, Golden SS, Hardin PE, Thomas TL, Zoran MJ. 2005 Circadian rhythms from multiple oscillators: lessons from diverse organisms. Nat Rev Genet 6, 544–556. (doi:10.1038/nrg1633)

3. Yerushalmi S, Green RM. 2009 Evidence for the adaptive significance of circadian rhythms. Ecol Lett 12, 970–981. (doi:10.1111/j.1461-0248.2009.01343.x)

4. Kondo T, Strayert CA, Kulkarnit RD, Taylor W, Ishiura M, Goldent SS, Hirschie C, Ii J. 1993 Circadian rhythms in prokaryotes: Luciferase as a reporter of circadian gene expression in cyanobacteria. Proc Natl Acad Sci U S A 90, 5672–5676.

5. Takahashi JS. 1995 Molecular neurobiology and genetics of circadian rhythms in mammals. Annu Rev Neurosci 18, 531–553. (doi:10.1146/annurev.ne.18.030195.002531)

6. King DP, Takahashi JS. 2000 Molecular genetics of circadian rhythms in mammals. Annu Rev Neurosci 23, 713–742.

7. Reppert SM, Weaver DR. 2001 Molecular analysis of mammalian circadian rhythms. Annual Review of Physiology 63, 647–676. (doi:10.1146/ANNUREV.PHYSIOL.63.1.647)

8. Yerushalmi S, Yakir E, Green RM. 2011 Circadian clocks and adaptation in Arabidopsis. Mol Ecol 20, 1155–1165. (doi:10.1111/j.1365-294X.2010.04962.x)

9. Rubin MJ, Brock MT, Baker RL, Wilcox S, Anderson K, Davis SJ, Weinig C. 2018 Circadian rhythms are associated with shoot architecture in natural settings. New Phytologist 219, 246–258. (doi:10.1111/nph.15162)

10. Rubin MJ et al. 2017 Circadian rhythms vary over the growing season and correlate with fitness components. Mol Ecol 26, 5528–5540. (doi:10.1111/mec.14287)

11. Koritala BSC et al. 2020 Habitat-specific clock variation and its consequence on reproductive fitness. J Biol Rhythms 35, 134–144.

12. Bloch G, Robinson GE. 2001 Reversal of honeybee behavioural rhythms. Nature 410, 1048–1048. (doi:10.1038/35074183)

13. Shemesh Y, Cohen M, Bloch G. 2007 Natural plasticity in circadian rhythms is mediated by reorganization in the molecular clockwork in honeybees. The FASEB Journal 21, 2304–2311. (doi:10.1096/FJ.06-8032COM)

14. Tessmar-Raible K, Raible F, Arboleda E. 2011 Another place, another timer: marine species and the rhythms of life. BioEssays 33, 165–172. (doi:10.1002/bies.201000096)

15. Barnwell FH. 1966 Daily and tidal patterns of activity in individual fiddler crab (Genus *Uca*) from the Woods Hole region. Biol Bull 130, 1–17.

16. Chabot CC, Ramberg-Pihl NC, Watson WH. 2016 Circalunidian clocks control tidal rhythms of locomotion in the American horseshoe crab, *Limulus polyphemus*. Mar Freshw Behav Physiol 49, 75–91. (doi:10.1080/10236244.2015.1127679)

17. Satoh A, Yoshioka E, Numata H. 2008 Circatidal activity rhythm in the mangrove cricket *Apteronemobius asahinai*. Biol Lett 4, 233–236. (doi:10.1098/rsbl.2008.0036)

18. Takekata H, Matsuura Y, Goto SG, Satoh A, Numata H. 2012 RNAi of the circadian clock gene *period* disrupts the circadian rhythm but not the circatidal rhythm in the mangrove cricket. Biol Lett 8, 488–491. (doi:10.1098/rsbl.2012.0079)

19. Takekata H, Numata H, Shiga S, Goto SG. 2014 Silencing the circadian clock gene *Clock* using RNAi reveals dissociation of the circatidal clock from the circadian clock in the mangrove cricket. J Insect Physiol 68, 16–22. (doi:10.1016/j.jinsphys.2014.06.012)

20. O’Neill JS, Lee KD, Zhang L, Feeney K, Webster SG, Blades MJ, Kyriacou CP, Hastings MH, Wilcockson DC. 2015 Metabolic molecular markers of the tidal clock in the marine crustacean *Eurydice pulchra*. Current Biology 25, R326–R327. (doi:10.1016/J.CUB.2015.02.052)

21. Mat AM, Massabuau JC, Ciret P, Tran D. 2014 Looking for the clock mechanism responsible for circatidal behavior in the oyster *Crassostrea gigas*. Mar Biol 161, 89–99. (doi:10.1007/s00227-013-2317-2)

22. Schnytzer Y, Simon-Blecher N, Li J, Ben-Asher HW, Salmon-Divon M, Achituv Y, Hughes ME, Levy O. 2018 Tidal and diel orchestration of behaviour and gene expression in an intertidal mollusc. Sci Rep 8. (doi:10.1038/s41598-018-23167-y)

23. Mat AM, Perrigault M, Massabuau JC, Tran D. 2016 Role and expression of *cry1* in the adductor muscle of the oyster *Crassostrea gigas* during daily and tidal valve activity rhythms. Chronobiol Int 33, 949–963. (doi:10.1080/07420528.2016.1181645)

24. Tran D, Perrigault M, Ciret P, Payton L. 2020 Bivalve mollusc circadian clock genes can run at tidal frequency. Proceedings of the Royal Society B: Biological Sciences 287. (doi:10.1098/rspb.2019.2440)

25. Kwiatkowski ER, Schnytzer Y, Rosenthal JJC, Emery P. 2023 Behavioral circatidal rhythms require *Bmal1* in *Parhyale hawaiensis*. Current Biology 33, 1867–1882. (doi:10.1016/J.CUB.2023.03.015)

26. Yokomizo T, Takahashi Y. 2022 Endogenous rhythm variation and adaptation to the tidal environment in the freshwater snail, *Semisulcospira reiniana*. Front Ecol Evol 10, 1218. (doi:10.3389/FEVO.2022.1078234)

27. Yamanaka O, Takeuchi R. 2018 UMATracker: An intuitive image-based tracking platform. Journal of Experimental Biology 221, 1–5. (doi:10.1242/jeb.182469)

28. Bolger AM, Lohse M, Usadel B. 2014 Trimmomatic: A flexible trimmer for Illumina sequence data. Bioinformatics 30, 2114–2120. (doi:10.1093/bioinformatics/btu170)

29. Grabherr MG et al. 2011 Full-length transcriptome assembly from RNA-Seq data without a reference genome. Nat Biotechnol 29, 644–652. (doi:10.1038/nbt.1883)

30. Li B, Dewey CN. 2014 RSEM: accurate transcript quantification from RNA-seq data with or without a reference genome. BMC Bioinformatics 12. (doi:10.1201/b16589)

31. Peñaloza C, Gutierrez AP, Eöry L, Wang S, Guo X, Archibald AL, Bean TP, Houston RD. 2021 A chromosome-level genome assembly for the Pacific oyster *Crassostrea gigas*. Gigascience 10, 1–9. (doi:10.1093/GIGASCIENCE/GIAB020)

32. Schmid B, Helfrich-Förster C, Yoshii T. 2011 A new ImageJ plug-in ‘actogramJ’ for chronobiological analyses. J Biol Rhythms 26, 464–467. (doi:10.1177/0748730411414264)

33. Thaben PF, Westermark PO. 2014 Detecting rhythms in time series with rain. J Biol Rhythms 29, 391–400. (doi:10.1177/0748730414553029)

34. Kuintzle RC, Chow ES, Westby TN, Gvakharia BO, Giebultowicz JM, Hendrix DA. 2017 Circadian deep sequencing reveals stress-response genes that adopt robust rhythmic expression during aging. Nat Commun 8. (doi:10.1038/ncomms14529)

35. Yu G, Wang LG, Han Y, He QY. 2012 ClusterProfiler: An R package for comparing biological themes among gene clusters. OMICS 16, 284–287. (doi:org/10.1089/omi.2011.0118)

36. Sharma VK. 2003 Adaptive significance of circadian clocks. Chronobiol Int 20, 901–919. (doi:10.1081/CBI-120026099)

37. Urabe M. 1998 Diel change of activity and movement on natural river beds in *Semisulcospira reiniana*. The Japanese journal of malacology 57, 17–27. (doi:10.18941/venusjjm.57.1_17)

38. Beaulé C. 2008 Aschoff’s Rules. Encyclopedia of Neuroscience, 190–193. (doi:10.1007/978-3-540-29678-2_383)

39. Hughes ME, DiTacchio L, Hayes KR, Vollmers C, Pulivarthy S, Baggs JE, Panda S, Hogenesch JB. 2009 Harmonics of circadian gene transcription in mammals. PLoS Genet 5. (doi:10.1371/journal.pgen.1000442)

40. Zhu B. 2020 Decoding the function and regulation of the mammalian 12-h clock. J Mol Cell Biol 12, 752–758. (doi:10.1093/jmcb/mjaa021)

41. Zhu B, Zhang Q, Pan Y, Mace EM, York B, Antoulas AC, Dacso CC, O’Malley BW. 2017 A cell-autonomous mammalian 12 hr clock coordinates metabolic and stress rhythms. Cell Metab 25, 1305–1319. (doi:10.1016/j.cmet.2017.05.004)

42. Zhang L, Hastings MH, Green EW, Tauber E, Sladek M, Webster SG, Kyriacou CP, Wilcockson DC. 2013 Dissociation of circadian and circatidal timekeeping in the marine crustacean *Eurydice pulchra*. Current Biology 23, 1863–1873. (doi:10.1016/j.cub.2013.08.038)

43. Satoh A, Terai Y. 2019 Circatidal gene expression in the mangrove cricket *Apteronemobius asahinai*. Sci Rep 9. (doi:10.1038/s41598-019-40197-2)

44. Mat AM et al. 2020 Biological rhythms in the deep-sea hydrothermal mussel *Bathymodiolus azoricus*. Nat Commun 11. (doi:10.1038/s41467-020-17284-4)

45. Hastings MH, Naylor E. 1980 Ontogeny of an endogenous rhythm in *Eurydice pulchra*. J Exp Mar Biol Ecol 46, 137–145. (doi:10.1016/0022-0981(80)90027-1)

46. Palmer JD. 1995 Review of the dual-clock control of tidal rhythms and the hypothesis that the same clock governs both circatidal and circadian rhythms. Chronobiol Int 12, 299–310.

47. Enright JT. 1976 Plasticity in an isopod’s clockworks: shaking shapes form and affects phase and frequency. Journal of Comparative Physiology A 107, 13–37. (doi:10.1007/BF00663916)

