## Supplementary Information for "Plasticity of circadian and circatidal rhythms in activity and transcriptomic dynamics in a freshwater snail"

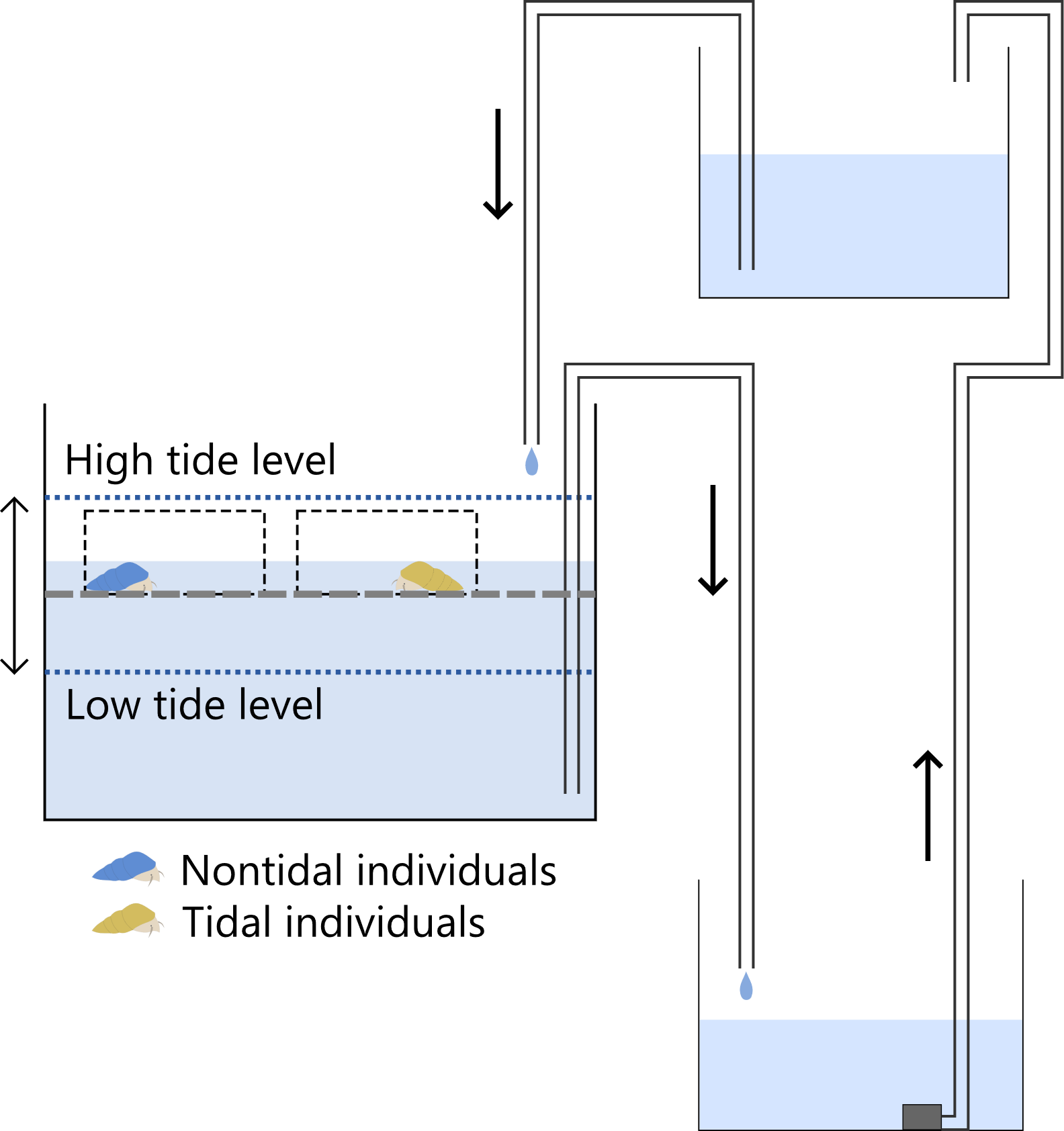

**Figure S1.** The structure of tidal simulation system. Snails were placed in the central tank (Main tank) and entrained to the tidal cycle. Water was supplied to and drained from the main tank using siphons and then pumped from the lower tank to the upper tank. In the main tank, snails were placed in the containers at the height of the dashed line. The dotted lines represent water levels of the high and low tides.

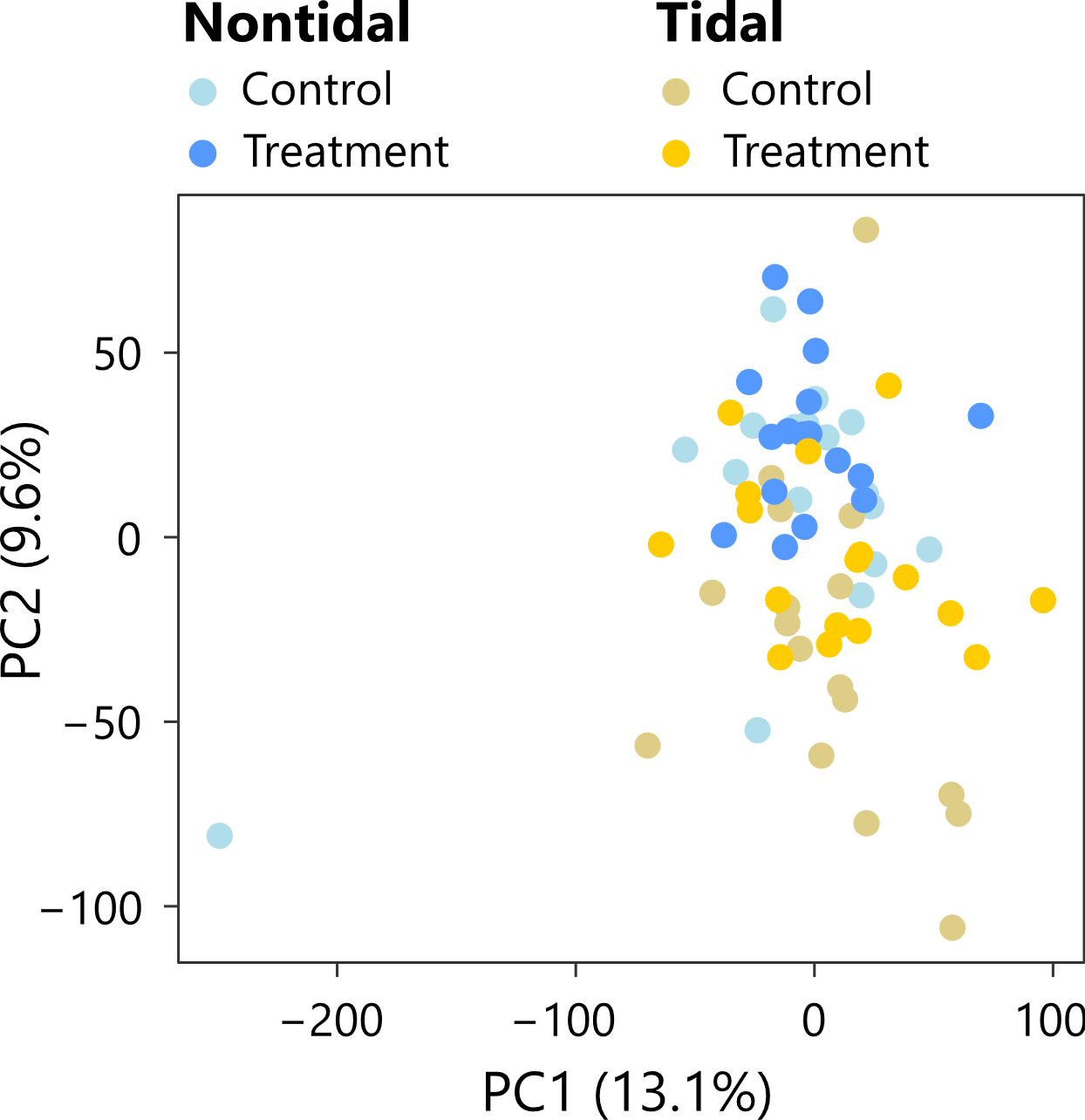

**Figure S2.** Probabilistic PCA of the expression of all genes after the filtering in nontidal and tidal populations using all samples.

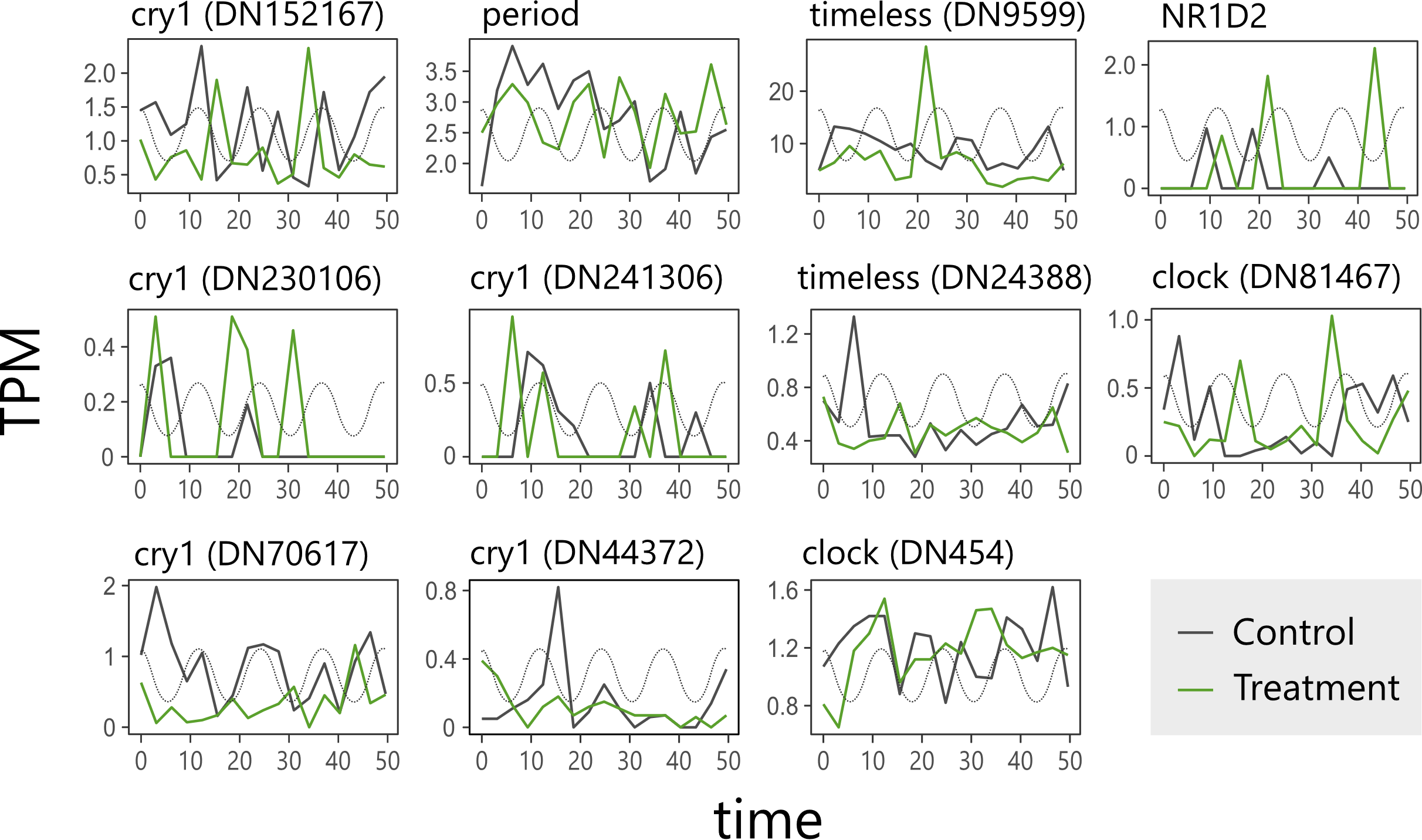

**Figure S3.** The expression patterns of circadian clock genes of the control (gray) and treatment (green) groups of the nontidal population under the DD condition. The simulated tidal cycle is shown as a gray dotted line.

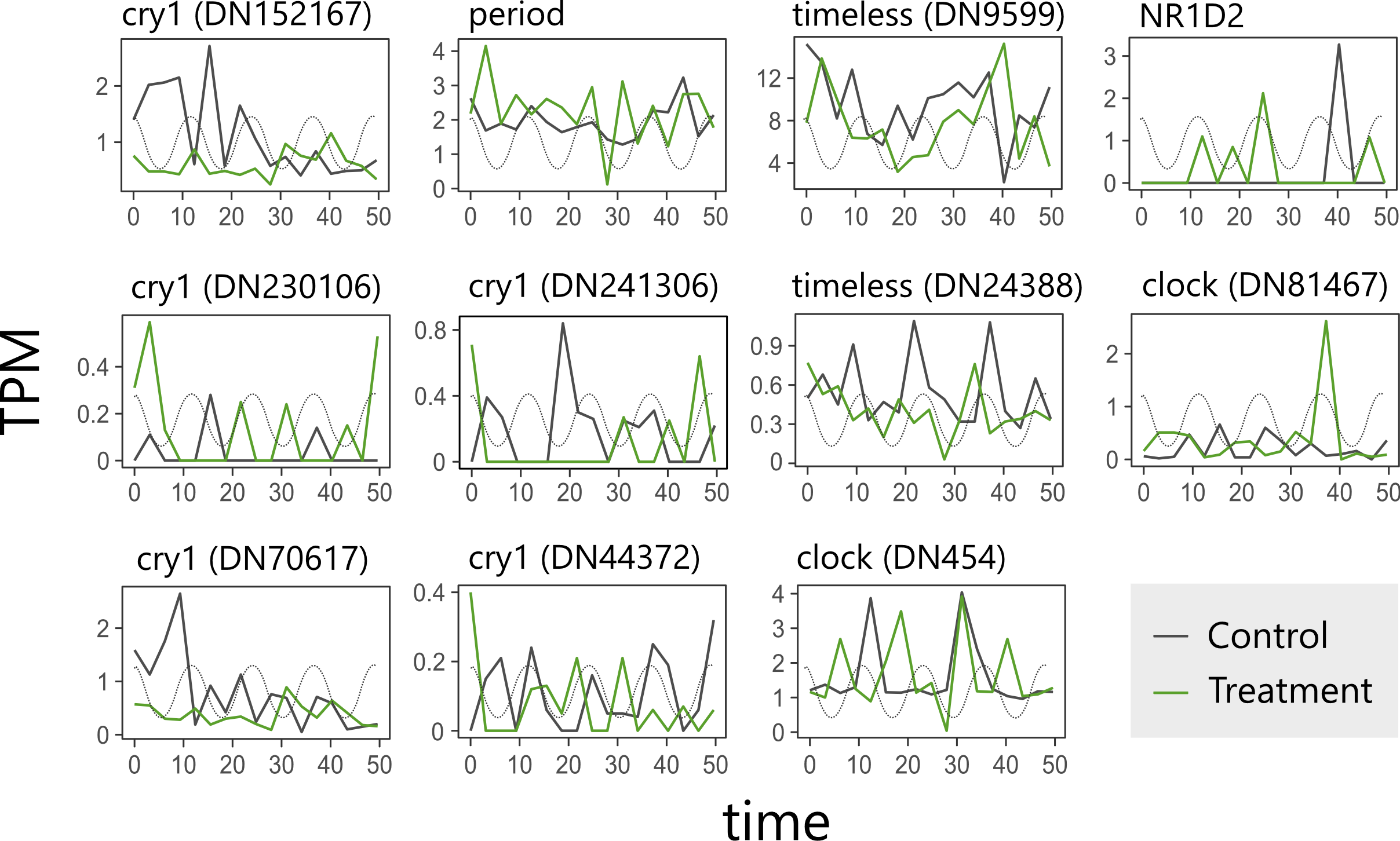

**Figure S4.** The expression patterns of circadian clock genes of the control (gray) and treatment (green) groups of the tidal population under the DD condition. The simulated tidal cycle is shown as a gray dotted line.

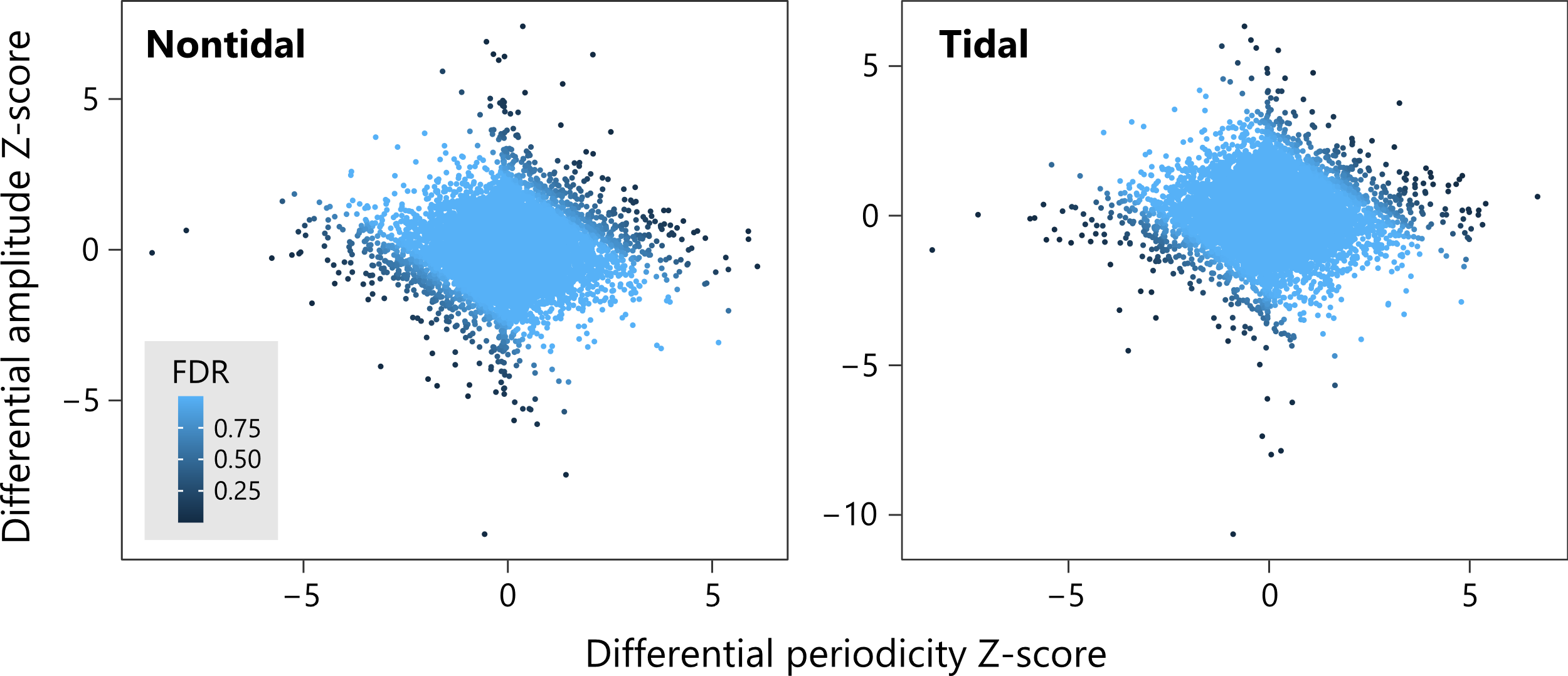

**Figure S5.** Differentiation of circatidal rhythmicity in the transcriptome between the control and treatment groups. Genes with higher differential periodicity Z-score (horizontal axis) exhibit stronger cyclical oscillation patterns of circatidal periods in the treatment group. Genes with a higher differential amplitude Z-score (vertical axis) show larger differences between their expression peak and trough in the treatment group. Genes are colored based on their false discovery rate (FDR) of *S*_DR_. Genes with positive and negative *S*_DR_ show increased and decreased rhythmicity in the treatment group, respectively.

**Table S1.** Genes that significantly increased rhythmicity of the circatidal period in the treatment group of the nontidal population. FDR is computed for *S*_DR_ using a Gaussian distribution based on the fit to the empirical distribution.

| Transcripts | *S*_DR_ | FDR | Description |
| --- | --- | --- | --- |
| DN21673_c2_g1 | 6.03 | 3.0 × 10^−6^ | transcription elongation factor 1 homolog isoform X2 |
| DN192135_c0_g1 | 5.48 | 6.7 × 10^−5^ | TIP41-like protein isoform X2 |
| DN449_c2_g1 | 4.82 | 0.0015 | uncharacterized protein LOC117689363 |
| DN105095_c2_g2 | 4.58 | 0.0039 | uncharacterized protein LOC117692043 |
| DN207785_c0_g1 | 4.53 | 0.0045 | uncharacterized protein LOC105346071 isoform X3 |
| DN39152_c0_g1 | 4.49 | 0.0050 | neurogenic locus notch homolog protein 2 isoform X17 |
| DN2536_c14_g1 | 4.46 | 0.0054 | sorting nexin-14 isoform X1 |
| DN18313_c0_g1 | 4.40 | 0.0058 | uncharacterized protein LOC117691072 isoform X1 |
| DN2970_c2_g2 | 4.32 | 0.0079 | tRNA (adenine(58)-N(1))-methyltransferase non-catalytic subunit TRM6 |
| DN6087_c0_g1 | 4.27 | 0.0090 | kinesin-like protein KIF21A isoform X14 |
| DN53630_c0_g1 | 3.97 | 0.030 | ABC transporter F family member 4 isoform X2 |
| DN93751_c0_g1 | 3.92 | 0.036 | multiple epidermal growth factor-like domains protein 10 isoform X3 |
| DN30054_c0_g1 | 3.83 | 0.045 | dual adapter for phosphotyrosine and 3-phosphotyrosine and 3-phosphoinositide isoform X3 |

**Table S2.** Genes that significantly increased rhythmicity of the circatidal period in the treatment group of the tidal population. FDR is computed for *S*_DR_ using a Gaussian distribution based on the fit to the empirical distribution.

| Transcripts | *S*_DR_ | FDR | Description |
| --- | --- | --- | --- |
| DN19687_c0_g1 | 5.16 | 2.9 × 10^−4^ | LOW QUALITY PROTEIN: ribonuclease P protein subunit p29 |
| DN50332_c0_g1 | 4.94 | 7.5 × 10^−4^ | uncharacterized protein LOC105334930 isoform X6 |
| DN6135_c0_g1 | 4.33 | 0.0098 | uncharacterized protein LOC105332169 isoform X3 |
| DN702_c1_g1 | 4.19 | 0.015 | glutaminyl-peptide cyclotransferase |
| DN7_c3_g1 | 4.13 | 0.018 | uncharacterized protein LOC117688477 |
| DN10387_c0_g1 | 4.08 | 0.020 | S-adenosylmethionine synthase |
| DN210683_c0_g1 | 4.05 | 0.021 | solute carrier family 35 member F5 isoform X3 |
| DN148503_c0_g1 | 4.02 | 0.022 | CD209 antigen-like protein E |
| DN48586_c2_g1 | 4.02 | 0.022 | uncharacterized protein LOC117692043 |
| DN140387_c0_g1 | 3.87 | 0.036 | uncharacterized protein LOC117681374 isoform X1 |
| DN67461_c2_g1 | 3.86 | 0.037 | zinc finger protein 665 |
| DN312_c13_g1 | 3.84 | 0.038 | uncharacterized protein LOC105333729 isoform X3 |
| DN177547_c2_g1 | 3.81 | 0.040 | uncharacterized protein LOC117685077 |
| DN6762_c5_g1 | 3.81 | 0.040 | uncharacterized protein LOC105341978 |
| DN618_c4_g2 | 3.80 | 0.041 | uncharacterized protein LOC117681191 isoform X1 |
| DN7859_c1_g1 | 3.75 | 0.047 | transcription elongation factor A N-terminal and central domain-containing protein 2-like |
